# Plasma after both SARS-CoV-2 boosted vaccination and COVID-19 potently neutralizes BQ.1.1 and XBB.1

**DOI:** 10.1101/2022.11.25.517977

**Authors:** David J Sullivan, Massimo Franchini, Jonathon W. Senefeld, Michael J. Joyner, Arturo Casadevall, Daniele Focosi

**Affiliations:** Johns Hopkins Bloomberg School of Public Health and School of Medicine, Baltimore, MD 21218, USA; Division of Transfusion Medicine, Carlo Poma Hospital, 46100 Mantua, Italy; Department of Anesthesiology & Perioperative Medicine, Mayo Clinic, Rochester, MN 55902, USA; North-Western Tuscany Blood Bank, Pisa University Hospital, 56124 Pisa, Italy

**Keywords:** convalescent plasma, SARS-CoV-2, COVID-19, BQ.1.1, XBB, BF.7: virus neutralization

## Abstract

**Objectives:** Recent 2022 SARS-CoV-2 Omicron variants, have acquired resistance to most neutralizing anti-Spike monoclonal antibodies authorized, and the BQ.1.* sublineages are notably resistant to all authorized monoclonal antibodies. Polyclonal antibodies from individuals both vaccinated and recently recovered from Omicron COVID-19 (VaxCCP) could retain new Omicron neutralizing activity.

**Methods:** Here we reviewed BQ.1.* virus neutralization data from 920 individual patient samples from 43 separate cohorts defined by boosted vaccinations with or without recent Omicron COVID-19, as well as infection without vaccination.

**Results:** More than 90% of the plasma samples from individuals in the recently (within 6 months) boosted VaxCCP study cohorts neutralized BQ.1.1, and BF.7 with 100% neutralization of WA-1, BA.4/5, BA.4.6 and BA.2.75. The geometric mean of the geometric mean 50% neutralizing titers (GM (GMT_50_) were 314, 78 and 204 for BQ.1.1, XBB.1 and BF.7, respectively. Compared to VaxCCP, plasma sampled from COVID-19 naïve subjects who also recently within 6 months received at least a third vaccine dose had about half of the GM (GMT_50_) for all viral variants.

**Conclusions:** Boosted VaxCCP characterized by either recent vaccine dose or infection event within 6 months represents a robust, variant-resilient, passive immunotherapy against the new Omicron BQ.1.1, XBB.1 and BF.7 variants.

## Introduction

In immunocompromised (IC) patients both passive immunotherapies and small molecule antivirals are often necessary to treat COVID-19 or eliminate persistently high SARS-CoV-2 viral load. Chronic, persistent viral loads increase both transmission and mutation risk, and prevent administration of the required immunosuppressive/antineoplastic therapies(1). Small molecule antivirals have not been formally validated for IC patients, who often have contraindications, and the convergent evolution of the Omicron variant of concern (VOC) has led to inefficacy of all the anti-Spike monoclonal antibodies (mAbs) authorized so far for both treatment or prevention, e.g. in the highly prevalent BQ.1.* sublineages(2). The other rapidly growing XBB.* and BF.7 sublineages are also highly resistant to anti-Spike mAbs(3). Polyclonal plasma from individuals who are both vaccinated and had COVID-19 (VaxCCP) has more than ten times the antibody levels capable of neutralizing pre-Omicron variants as well as Omicron variants BA.1 through BA.4/5(4, 5). Polyclonal COVID-19 convalescent plasma (CCP) has thousands of distinct antibody specificities of different isotypes, including many capable of SARS-CoV-2 neutralization. High-titer pre-Omicron CCP contains Omicron neutralizing activity despite being collected before variant appearance(4, 5).

Given that CCP remains a recommended therapy for IC(1, 6, 7), we systematically reviewed recent primary research for neutralization results against BQ.1.1 by plasma collected from vaccinated subjects with or without COVID-19 or after recent Omicron infection alone.

## Results

Ten articles were included (**Figure 1**) which contained virus neutralizations with WA-1, BQ.1.1, BA.4/5, BA.4.6, XBB.1 and BF.7, assessed with either live authentic SARS-CoV-2 or SARS-CoV-2 pseudovirus neutralization assays and represented data from 920 patients (**Supplementary Table 1**). Qu *et al*. in the USA reported on Spring and Summer 2022 breakthrough infections with BA.1 and BA.4/5 in two sampled cohorts with predominantly unvaccinated individuals, as well as a third cohort of healthcare workers after a single monovalent booster vaccination in the Fall of 2021(8) (**Table 1**). Zou *et al*. in the USA in the Summer and Fall of 2022 sampled individuals who had already received 3 mRNA BNT162b2 vaccinations with or without previous COVID-19, both before and about 4 weeks after a 4^th^ monovalent or bivalent vaccine booster vaccination(9). Miller *et al*. also in the USA sampled both before the 3^rd^ vaccination dose and about 4 weeks after monovalent mRNA vaccination in the Fall of 2021, as well as with the 4th vaccine dose in the Summer or Fall of 2022, with either monovalent or bivalent booster vaccinations in Fall of 2022 in those with no documented COVID-19(10). Cao *et al*. in China investigated BQ.1.1 neutralizations from plasma of 4 cohorts after 3 doses of CoronaVac (Fall 2021) without COVID-19 or 2-12 weeks after BA.1, BA.2 and BA.5 infection(3). Planas *et al*. in France evaluated GMT_50_in plasma from individuals both 4 and 16 weeks after a third monovalent mRNA vaccine dose in the Fall of 2021 as well as 12 and 32 weeks after vaccine breakthrough BA.1/2 or BA.5 infection(11). Davis *et al* in the USA sampled after the 3^rd^ mRNA vaccine monovalent dose in the Fall of 2021 and also after either a 4th monovalent mRNA dose or a bivalent (wild-type + BA.4/5) vaccine dose in the Summer and Fall of 2022(12). Kurhade *et al* in the USA also compared GMT_50_after the 4^th^ monovalent vaccine dose or 3 mRNA doses with the 4^th^ the bivalent dose without COVID-19 and also after bivalent boost with recent COVID-19(13). Wang *et al* in the USA compared GMT_50_ after three vaccine doses, the 4^th^ monovalent vaccine dose or 3 mRNA doses with the 4^th^ the bivalent dose without COVID-19, and also after 2-3 vaccine doses and recent BA.2 breakthrough infection or 3-4 mRNA vaccine doses and recent BA.4/5 breakthrough infection(14). Ito et al in Japan compared breakthrough infections after BA.2 and BA.5 after 2-3 doses of mRNA vaccines in the Spring and Summer of 2022(15). Akerman et al in Australia characterized neutralizing antibodies in four groups 1) sampling one to three months after 3 doses of mRNA vaccines with an Omicron infection in 2022; 2) sampling 3 months after 4 doses of mRNA vaccine; 3) sampling 6 months after 3 doses of mRNA vaccine and 4) sampling 3-6 months after last vaccine in a larger cohort who had the original WA-1 infection in early 2020 as well as 3 more doses of mRNA vaccine(16).

**Figure 1.**
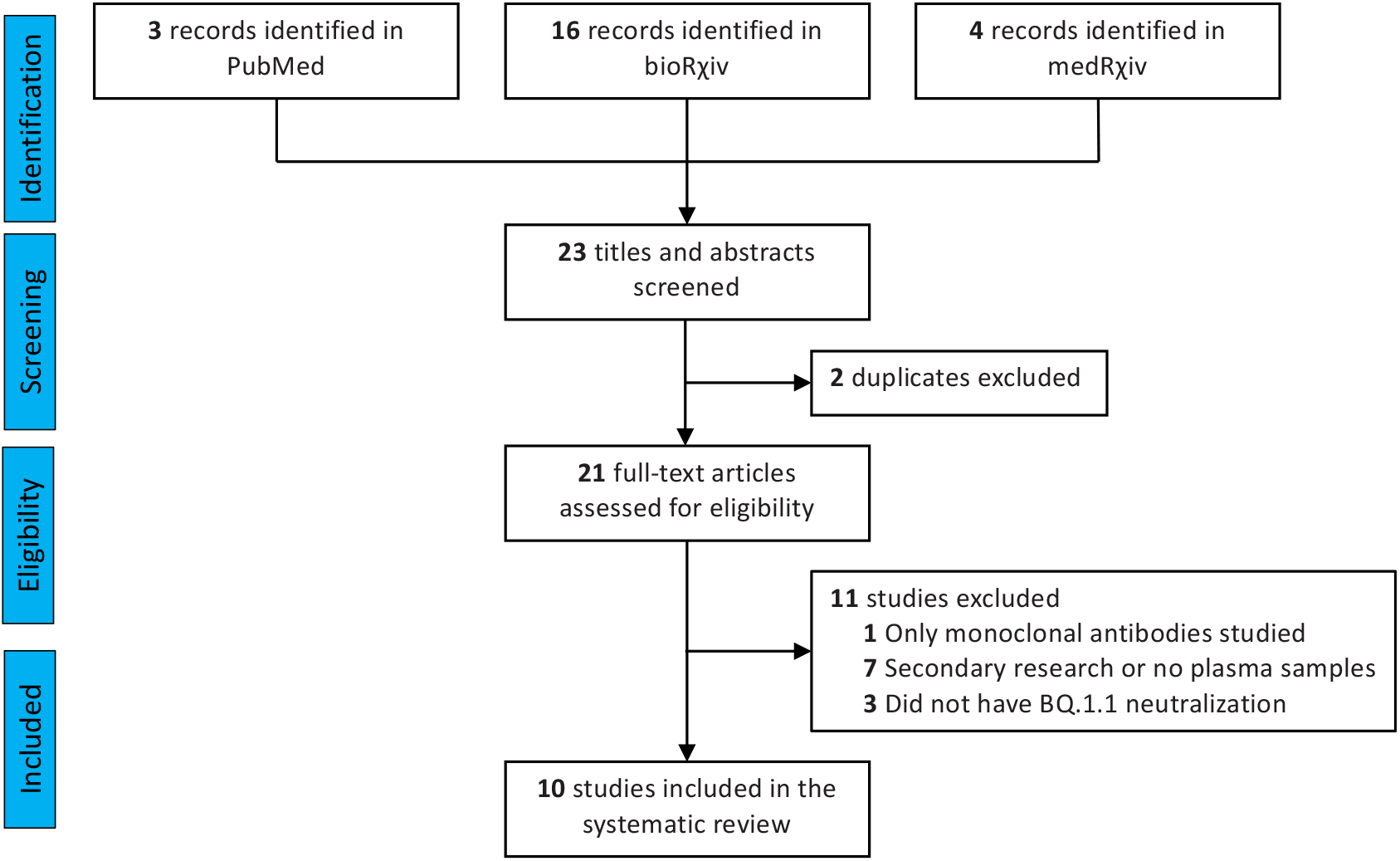
PRISMA flowchart for the current study. Number of records identified from various sources, excluded by manual screening, found eligible and included according to subgroup analyses.

**Table 1.** Synopsis of included studies, reporting plasma sources, epoch of sampling, region, time since vaccination/infection to plasma sampling, and sample size. The cohorts were split into three groups-1) boosted vaccinations and recent COVID-19 (VaxCCP), 2) boosted vaccines only without documented COVID-19 (Vac only) and 3) infection alone or pre-boosted sampling before 3^rd^ or 4^th^ vaccine dose (Infection only or pre-boost)

| Study | Vaccine and COVID-19 history at sample time | Group | Time period of plasma sampling | Geography | Sampling time mean or median (min-max) | Sample number |
| --- | --- | --- | --- | --- | --- | --- |
| Cao(3) | 3xCorVac+BA.1 inf | VaxCCP | Spring 2022 | China | 5-7 weeks post hosp admit (42 weeks avg) | 50 |
| Cao(3) | 3xCorVac +BA.2 inf | VaxCCP | Summer 2022 | China | 3-11 weeks post hosp admit (8 weeks mean) | 39 |
| Cao(3) | 3xCorVac +BA.5 inf | VaxCCP | Summer/Fall 2022 | China | 2-11 weeks (mean 5 weeks) | 36 |
| Zou(9) | 4xBNT162b2+BTI | VaxCCP | Summer/Fall 2022 | USA | 4 weeks post dose | 20 |
| Planas(11) | mRNAvac3+ BA.1/2 inf | VaxCCP | Spring/Fall 2022 | France | 32 weeks post BTI BA.1/2 | 13 |
| Wang(14) | 2-3xmRNAvac+ BA.2 BTI | VaxCCP | Spring/Fall 2022 | USA | 2-23 weeks (mean 6 wk (3over 90 days)) | 14 |
| Wang(14) | 3-4xmRNAvac+ BA.4/5 BTI | VaxCCP | Summer/Fall 2022 | USA | 2-8 weeks (mean 4 weeks) | 20 |
| Kurhade(13) | 3xmRNAvac+bivalent+BTI | VaxCCP | Fall 2022 | USA | 4 weeks post with infection history | 23 |
| Ito(15) | 2-3xmRNAvac+BA.2 BTI | VaxCCP | Spring 2022 | Japan | 2-8 weeks | 14 |
| Ito(15) | 2-3xmRNAvac+BA.5 BTI | VaxCCP | Summer 2022 | Japan | 2-3 weeks | 20 |
| Zou(9) | 3xBNT162b2+bivalent+BTI | VaxCCP | Summer/Fall 2022 | USA | 4 weeks post dose | 19 |
| Akerman(16) | 3xmRNA +bivalent | VaxCCP | Fall 2022 | Australia | 4-12 weeks post BTI | 29 |
| Planas(11) | 3xmRNAvac+ BA.1/2 inf | VaxCCP | Spring/Fall 2022 | France | 12 weeks post BTI BA.1/2 | 16 |
| Planas(11) | 3xmRNAvac+ BA.5 inf | VaxCCP | Fall 2022 | France | 8 weeks post BTI BA.5 | 15 |
| Davis(12) | 3xmRNAvac | Vac only | Fall 2021 | USA | 1-4 weeks post boost | 12 |
| Kurhade(13) | 4xmRNAvac | Vac only | Summer 2022 | USA | 4-12 weeks | 25 |
| Cao(3) | 3xCorVac | Vac only | Fall 2021 | China | 4 weeks | 40 |
| Zou(9) | 4xBNT162b2 | Vac only | Summer/Fall 2022 | USA | 4 weeks post dose | 20 |
| Planas(11) | 3xmRNAvac | Vac only | Winter 2021/2022 | France | 16 weeks post 3rd dose | 10 |
| Wang(14) | 3xmRNAvac | Vac only | Fall 2021 | USA | 2-12 weeks (mean 5 weeks) | 14 |
| Wang(14) | 3xmRNAvac+monovalent | Vac only | Summer/Fall 2022 | USA | 3-4 weeks | 19 |
| Wang(14) | 3xmRNAvac+bivalent | Vac only | Summer/Fall 2022 | USA | 3-4 weeks | 21 |
| Davis(12) | 3xmRNAvac+monovalent | Vac only | Summer/Fall 2022 | USA | 10-15 weeks post boost | 12 |
| Akerman(16) | 4xmRNA | Vac only | Fall 2022 | Australia | 12 weeks | 23 |
| Akerman(16) | 3xmRNAvac after 2020 WA-1 | Vac only | Summer/Fall 2022 | Australia | 3-6 months | 47 |
| Kurhade(13) | 3xmRNAvac+bivalent | Vac only | Fall 2022 | USA | 4 weeks post | 29 |
| Davis(12) | 3xmRNAvac+bivalent | Vac only | Summer/Fall 2022 | USA | 2-6 weeks post booster (8 no vacc. 10 no infection) | 12 |
| Qu(8) | 3xmRNAvac | Vac only | Fall 2021 | USA | 2-13 weeks | 15 |
| Zou(9) | 3xBNT162b2+bivalent | Vac only | Summer/Fall 2022 | USA | 4 week post dose | 18 |
| Planas(11) | 3xmRNAvac | Vac only | Fall/Winter 2021 | France | 4 weeks post 3rd dose | 18 |
| Miller(10) | 3xBNT162b2 | Vac only | Fall 2021 | USA | 2-4 weeks | 16 |
| Miller(10) | 3xmRNA+monovalent | Vac only | Spring/Fall 2022 | USA | 2-9 weeks | 18 |
| Miller(10) | 3xmRNA +bivalent | Vac only | Fall 2022 | USA | 2-3 weeks | 15 |
| Qu(8) | BA.4/5 inf (17-unvac) | Inf only | Summer 2022 | USA | not stated | 20 |
| Qu(8) | Hosp BA.1 (6-unvac;5-2xmRNAvac) | Inf only | Spring 2022 | USA | 1 week post hospitalization | 15 |
| Zou(9) | 3xBNT162b2+BTI | preboost with BNT162b | Summer/Fall 2022 | USA | preboost with BNT162b (6-11 months post last dose) | 20 |
| Zou(9) | 3xBNT162b2 +BTI | preboost with bivalent | Summer/Fall 2022 | USA | preboost with bivalent (6-11 months post last dose) | 19 |
| Zou(9) | 3xBNT162b2 | preboost with bivalent | Summer/Fall 2022 | USA | preboost with bivalent (6-11 months post last dose) | 18 |
| Zou(9) | 3xBNT162b2 | preboost with BNT162b | Summer/Fall 2022 | USA | preboost with BNT162b (6-11 months post last dose) | 20 |
| Akerman(16) | 3xmRNA | Preboost | Fall 2022 | Australia | preboost (6 months) | 47 |
| Miller(10) | 2xBNT162b2 | preboost with BNT162b | Fall 2021 | USA | preboost (6-11 months post last dose) | 16 |
| Miller(10) | 3xmRNA | preboost with bivalent | Fall 2022 | USA | preboost with bivalent (6-11 months post last dose) | 15 |
| Miller(10) | 3xmRNA | preboost with monovalent | Spring/Fall 2022 | USA | preboost with monovalent (6-11 months post last dose) | 18 |

These diverse cohorts were assembled into 3 groups, 1) plasma after both 2-4 vaccine doses and COVID-19 (VaxCCP); 2) plasma from subjects after administration of 3-4 vaccine doses (i.e. boosted), but either self-reported as COVID-19-*naïve* or anti-nucleocapsid negative; and 3) Omicron infection without vaccination (CCP) as well as participants sampled 6 to 11 months after previous vaccine dose and before the booster vaccination. Boosted VaxCCP neutralized BQ.1.1, XBB.1 and BF.7 with approximately 3 times the dilutional potency of the vaccine-only or 2-6 times CCP/pre-boost vaccination groups for all viral variants (**Table 2 and Figure 2**). Importantly, while there was a 19-fold reduction in neutralization by boosted VaxCCP against BQ.1.1 compared to WA-1, more than 90% of the boosted VaxCCP samples neutralized BQ.1.1 as well as XBB.1 and BF.7 (**Table 2 and Figure 2c**). Three cohorts within the boosted VaxCCP group were below at 90% neutralization with one sampled late, 8 months after BA.1/2 breakthrough infection(11) and the other two from a single study after BA.2 and BA.5 (**Supplementary Table 2 and 3**). Except for the GMT (GMT_50_) against XBB.1 at 78, the other viral variant neutralizations were in the same range as pre-Alpha CCP neutralizing WA-1 (i.e., 311)(4). By comparison the large randomized clinical trial which effectively reduced outpatient COVID-19 progression to hospitalizations had a GMT_50_ of 60 for WA-1 with pre-Alpha CCP(17). Boosted vaccinations at 3-4 doses without COVID-19, showed GM (GMT_50_) of 118 for BQ.1.1, with only 6 of 23 cohorts over 90% neutralizations, for 79% overall (i.e. 326 of 414 individuals). Four separate studies(8),(13),(12),(10) characterized BQ.1.1 virus neutralizations with plasma after the new bivalent (wild-type + BA.4/5) mRNA vaccine booster in the Fall of 2022, with 88% (103 of 117 samples) neutralization activity within 4 weeks of bivalent booster (**Supplementary Table 3**).

**Table 2.**
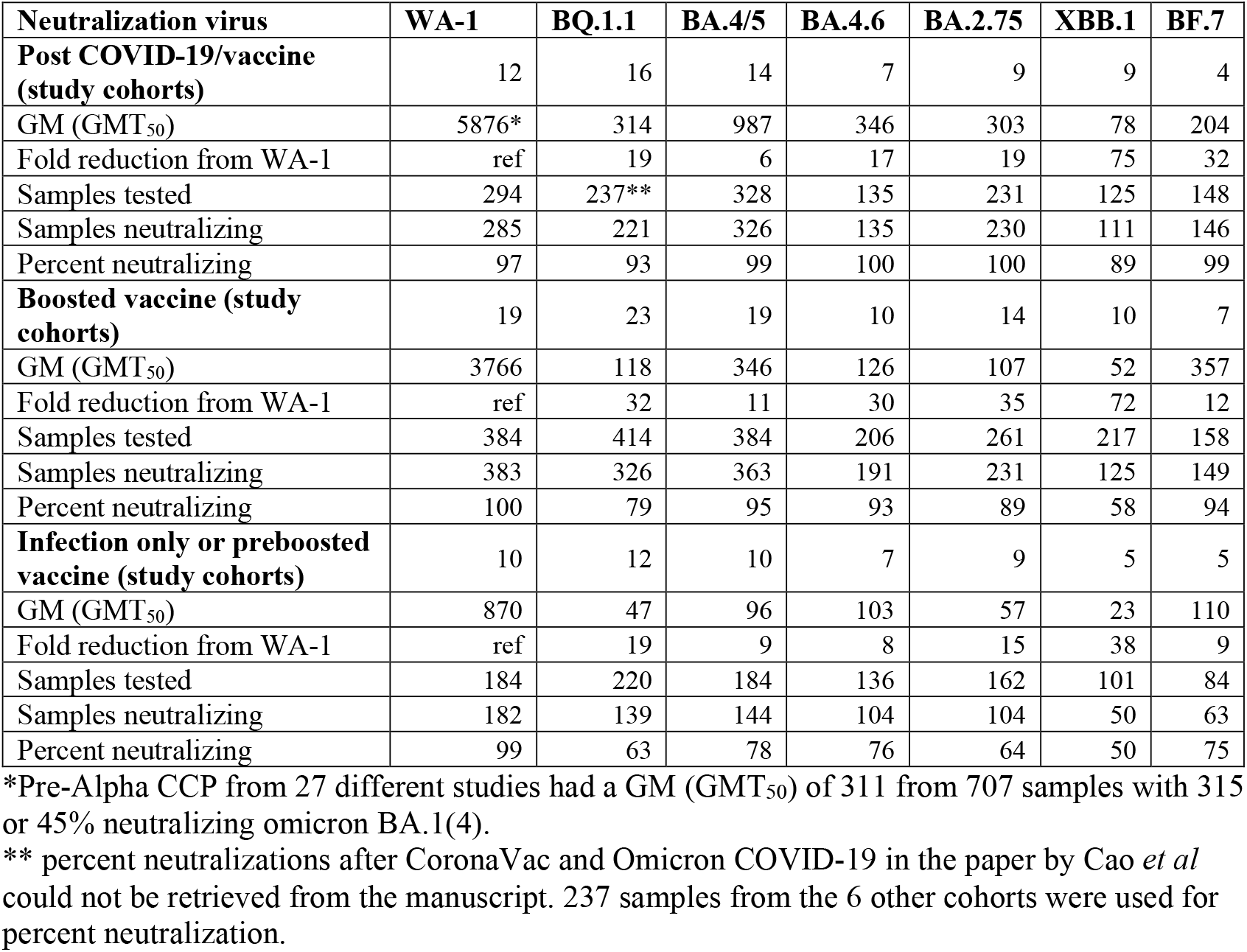
GM (GMT_50_) of plasma from three different sources against recent Omicron sublineages.

**Figure 2.**
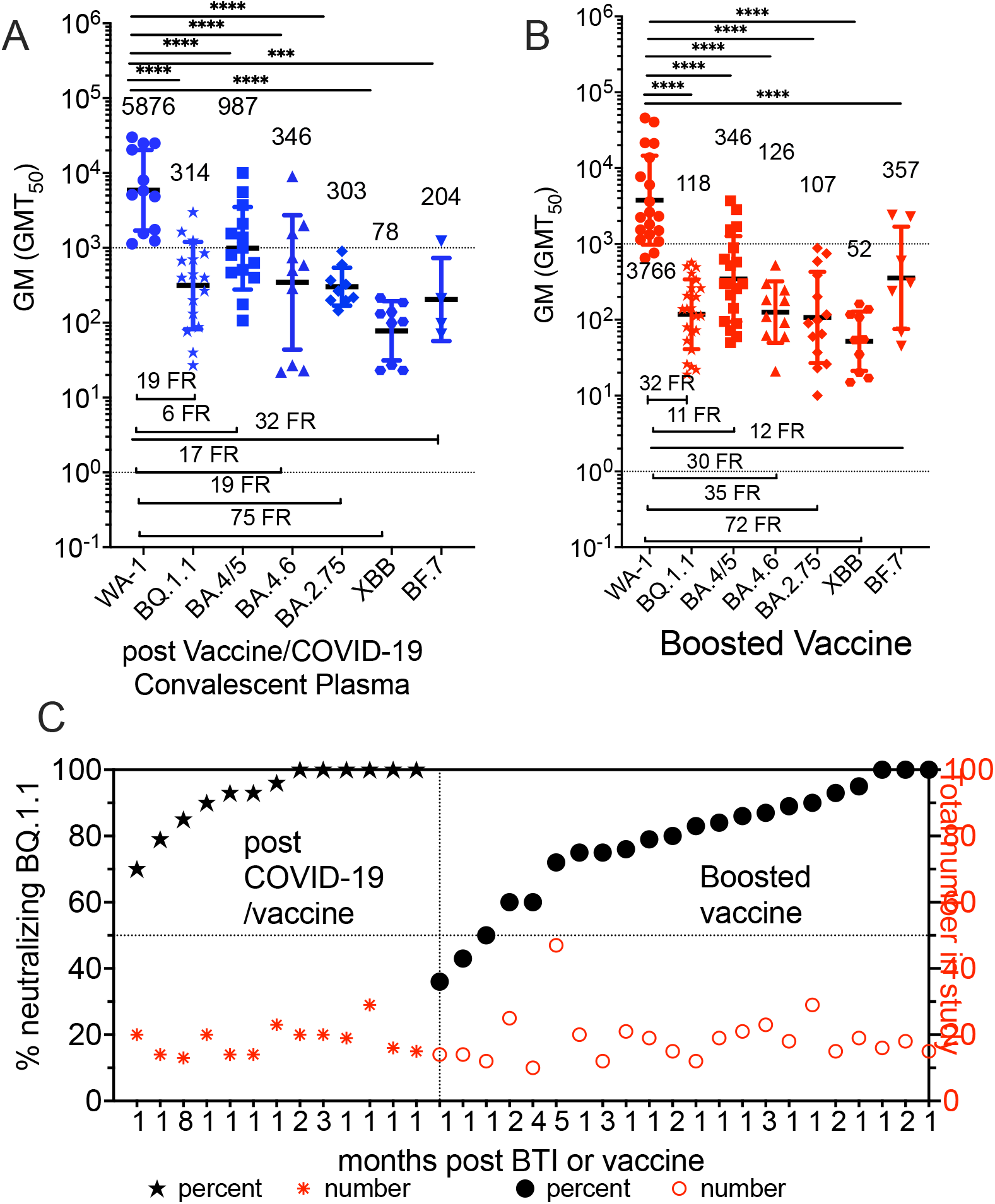
Neutralizing GMT (GMT_50_) against WA-1, BQ.1.1, BA.4/5, BA.4.6, BA.2.75, XBB, BF.7. A) post boosted vaccinations and COVID-19 and B) boosted vaccinated plasma without COVID-19. Geometric standard deviation for error bars, fold reduction (FR) below data, and number of studies above x-axis. Geomeans statistically significant in difference by multiple comparison in Tukey’s test are indicated. C) The percent of total samples within a study which neutralized Omicron BQ.1.1 graphed in increasing percentages on left y axis with the total number of samples tested on the right y axis.

Many studies performed virus neutralizations on samples drawn before the 3^rd^ or 4^th^ vaccine dose which were 6 to 11 months after last vaccine dose. The GM (GMT_50_)’s for BQ.1.1 and BA.2.75 were about 6 times reduced compared to VaxCCP even though the fold reductions were similar (**Figure 3, Table 2**). In agreement with lower GMT_50_ for neutralizations was the low percent neutralizing BQ.1.1 (63%), XBB.1 (50%), and BF.7 (75%) at 6 to 11 months after vaccination (**Figure 3, Table 2 and Supplementary Table 3**).

**Figure 3.**
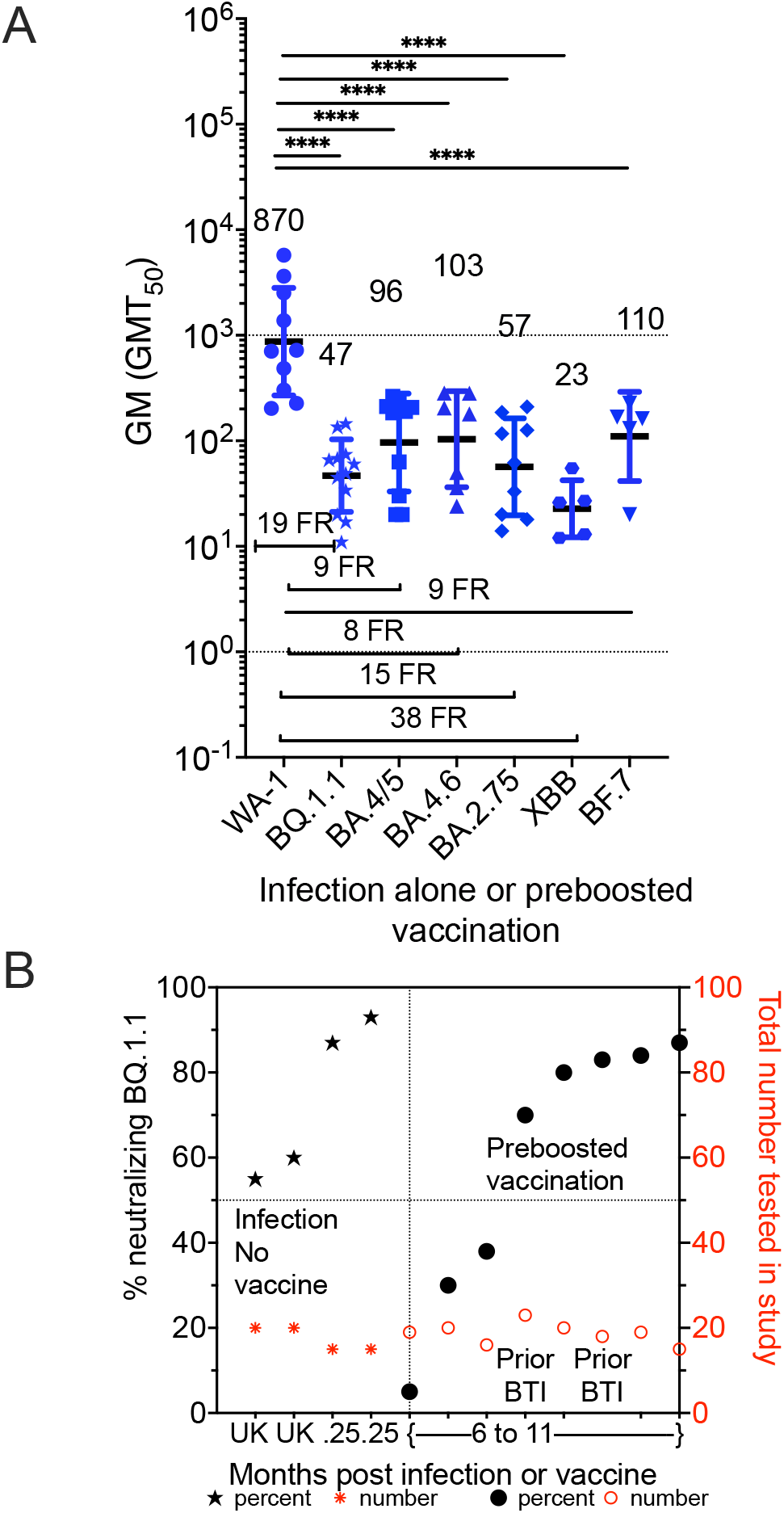
Geometric mean neutralizing titers (GMT_50_) against WA-1, BQ.1.1, BA.4/5, BA.4.6, BA.2.75, XBB, BF.7 A) plasma Omicron infection alone or pre-boosted-6 to 11 months after last vaccine dose sampled in 2021 or 2022. Geometric standard deviation for error bars, fold reduction (FR) above data, and number of studies above x-axis. GM (GMT_50_) statistically significant in difference by multiple comparison in Tukey’s test are indicated. B). The percent of total samples within a study which neutralized Omicron BQ.1.1 graphed in increasing percentages on left y-axis with the total number of samples tested on the right y-axis.

Five studies used the lentiviral pseudovirus assays, with diverse Spike proteins cloned in, while the other four were live virus assays using different cell types (**Supplementary Table 1**). Notably, Planas *et al* employed the IGROV-1 cell type for better growth of Omicron sublineages(11). While the single study fold reductions (FR) and percent neutralizations normalize the results between studies, the GMT_50_ can vary between studies even amongst the live authentic viral neutralization studies (e.g., mNeonGreen™ reporter assays versus cytopathic effects)(9, 13). We sorted the live authentic viral neutralizations from the pseudoviral neutralizations, plotting also the minimum and maximums (**Supplementary Figures 1-3**). In general, the live authentic SARS-CoV-2 neutralization assays for VaxCCP appeared to have similar antibody neutralization levels, with the single study by Cao et al(3) employing lentiviral pseudovirus with lower dilutional titers. In contrast, the GMT_50_ achieved with pseudoviral assays in the boosted vaccinations without COVID-19 appeared slightly higher than the ones achieved with authentic virus.

## Discussion

The FDA deemed CCP safe and effective for both immunocompetent and IC COVID-19 outpatients(6, 7, 18), and further extended its authorized use in the IC patient population in December 2021(7, 18), at a time when oral antiviral therapy promised a no transfusion outpatient solution and many anti-Spike mAbs were still effective.

Up until the present, CCP remained a backup bridge for IC patients, durable against the changing variants and as a salvage therapy in seronegative IC patients. With the recent advent of Omicron XBB.* and BQ.1.* defeating the remaining anti-Spike mAbs, boosted VaxCCP, recently collected within the last 6 months of either a vaccine dose or SARS-CoV-2 is likely to be the only viable remaining passive antibody therapy in the 2022-23 Winter for IC patients who have failed to make antibodies after vaccination and still require B-cell depleting drugs or immunosuppressive therapy. In a literature review of CCP from diverse VOC waves as well as boosted vaccinees and VaxCCP up to BA-1, VaxCCP showed higher neutralization titers against Omicron at levels above 300 dilutional GMT_50_^4^.

The accelerated evolution of SARS-CoV-2 VOCs has created the problem that the pharmaceutical development of additional mAbs is not worth the effort and cost given their expected short useful clinical life expectancy, so the anti-Spike mAb pipeline has remained stuck in 2022. High levels of antibodies in donor plasma from both boosted vaccinations and COVID-19 convalescent plasma (VaxCCP) neutralizing more than 93% of BQ.1.1and BF.7, with XBB.1 at 89%. Recently collected plasma within a 6 months window from those boosted vaccinees without prior documented COVID-19 had a 20-30% reduction in neutralization percent for BQ.1.1and XBB.1 with 10% reduction for the others and a third of the GM (GMT_50_) neutralizing antibody levels compared to VaxCCP. In those vaccinated with last dose more than 6 months prior to sample collection, both the neutralization percent and neutralizing antibody titers fell further, compared to the recently boosted VaxCCP group. Four studies (Planas(11), Zou(9), Cao(3) and Kurhade(13)) had directly comparative cohorts in the three groups which increases the robustness reduction in neutralizations with the vaccine only or more than 6 months to last vaccine or infection event compared to VaxCCP. The main limitation of our systematic review is the small number of studies reporting virus neutralization with BQ.1.1 with most available as pre-preprints without peer-review yet. However, we note that peer-review itself does not change GMT_50_ or neutralization numbers and the authors of these papers have considerable expertise in the topic.

Boosted VaxCCP has full potential to replace anti-Spike mAbs for passive antibody therapy of IC patients against recent Omicron sublineages, in the meanwhile polyclonal IgG formulations can be manufactured. VaxCCP qualification in the real-world will likely remain constrained on high-throughput serology, whose correlation with GMT_50_ is not perfect(19, 20). Nevertheless, the very high prevalence (93%) of Omicron-neutralizing antibodies and the high GM (GMT_50_) in recently boosted VaxCCP reassure about its potency, and further confirm that exact donor-recipient VOC matching is dispensable. Overall, our findings urge WHO to revise its guidelines and recommend boosted VaxCCP for therapy of COVID-19 in IC patients.

### Search strategy and selection criteria

On November 19, 2022 we initially searched PubMed, medRxiv and bioRxiv for manuscripts reporting BQ.1.1 neutralization, using English language as a restriction. Search of bioRxiv with same keywords now yields 17 records of which only 10 contained plasma viral neutralization data. Search of medRxiv produced 3 records which did not have BQ.1.1 neutralizations. PubMed retrieved 3 entries using (“BQ.1.1”) and (“neutralization”), one of which was focused on anti-Spike mAb alone(2) and the other 2 were duplicates from bioRxiv(8, 12). Articles underwent evaluation for data extraction by two assessors (DS and DF) with disagreements resolved by third assessor (AC). Articles lacking plasma BQ.1.1 virus neutralizations were excluded. The process of study selection is represented in the PRISMA flow diagram (**Figure 1**).

The type of viral assay (live or pseudovirus), time interval to blood sample, GMT_50_, minimum and maximum neutralizing 50% dilutional titer for WA-1 (pre-Alpha wild-type) and Omicron sublineages BQ.1.1, BA.4/5, BA.4.6, BA.2.75, XBB.1 and BF.7 and number out of total that neutralized Omicron were abstracted from study text, graphs and tables. Two studies (Wang(14) and Qu(8)) reported BQ.1 and those were separate cohorts in addition to BQ.1.1. Prism v. 9.4 (GraphPad Software, San Diego, CA, USA) was used for data analysis. While all manuscripts included neutralization data against WA-1, BQ.1.1, BA.4/5 and BA.2.75, only a subset of manuscripts included neutralization data for BA.4.6, XBB.1 and BF.7 which were assembled for relevance to present circulating variants. Historic early Omicron partial neutralization data on variants like BA.1 or BA.2 were excluded because of the full set data with BA.4/5 and BA.2.75.

Statistical significance between log_10_ transformed GMT_50_ was investigated using Tukey’s test. The multiple comparison test was a two-way ANOVA with Alpha 0.05 on log transformed GMT_50_. The log normal test was performed on WA-1, BQ.1.1, BA.4/5, BA.4.6, XBB.1 and BF.7 virus GMT_50_. Two studies(10, 11) reported the median titer rather than the GMT_50_. Compiled data abstracted from the published studies is available in the supplementary dataset.

## Abbreviations

VOC: variant of concern;
VaxCCP: plasma from both vaccinated and COVID-19 convalescent subjects

---

**Supplementary Table 1.** Virus neutralization assays.

| Reference | Assay | Virus | Replication-competent cells | neutralization threshold |
| --- | --- | --- | --- | --- |
| Qu(8) | Pseudovirus | Lentiviral pseudovirus | HEK293T-ACE-2 | 80 |
| Miller(10) | Pseudovirus | Lentiviral pseudovirus | HEK293T-ACE-2 | 20 |
| Cao(3) | Pseudovirus | VSV pseudovirus | HEK293T-ACE-2 | 20 |
| Wang(14) | Pseudovirus | VSV pseudovirus | VeroE6- | 100 |
| Ito(15) | Pseudovirus | Lentiviral pseudovirus | HEK293T-ACE-2 | 150 |
| Akerman(16) | Live virus | Live authentic SARS-CoV-2 | VeroE6-TMPRSS2 | 20 |
| Davis(12) | Live virus | Live authentic SARS-CoV-2 | VeroE6-TMPRSS2 | 20 |
| Kurhade(13) | Live virus | mNeonGreen reporter USA-WA1/2020 SARS-CoV-2 | Vero E6-TMPRSS2 | 20 |
| Planas(11) | Live virus | Live authentic SARS-CoV-2 | IGROV-1 or Vero E6-TMPRSS2 | 30 |
| Zou(9) | Live virus | mNeonGreen reporter USA-WA1/2020 SARS-CoV-2 | Vero E6-TMPRSS2 | 20 |

**Supplementary Table 2.**
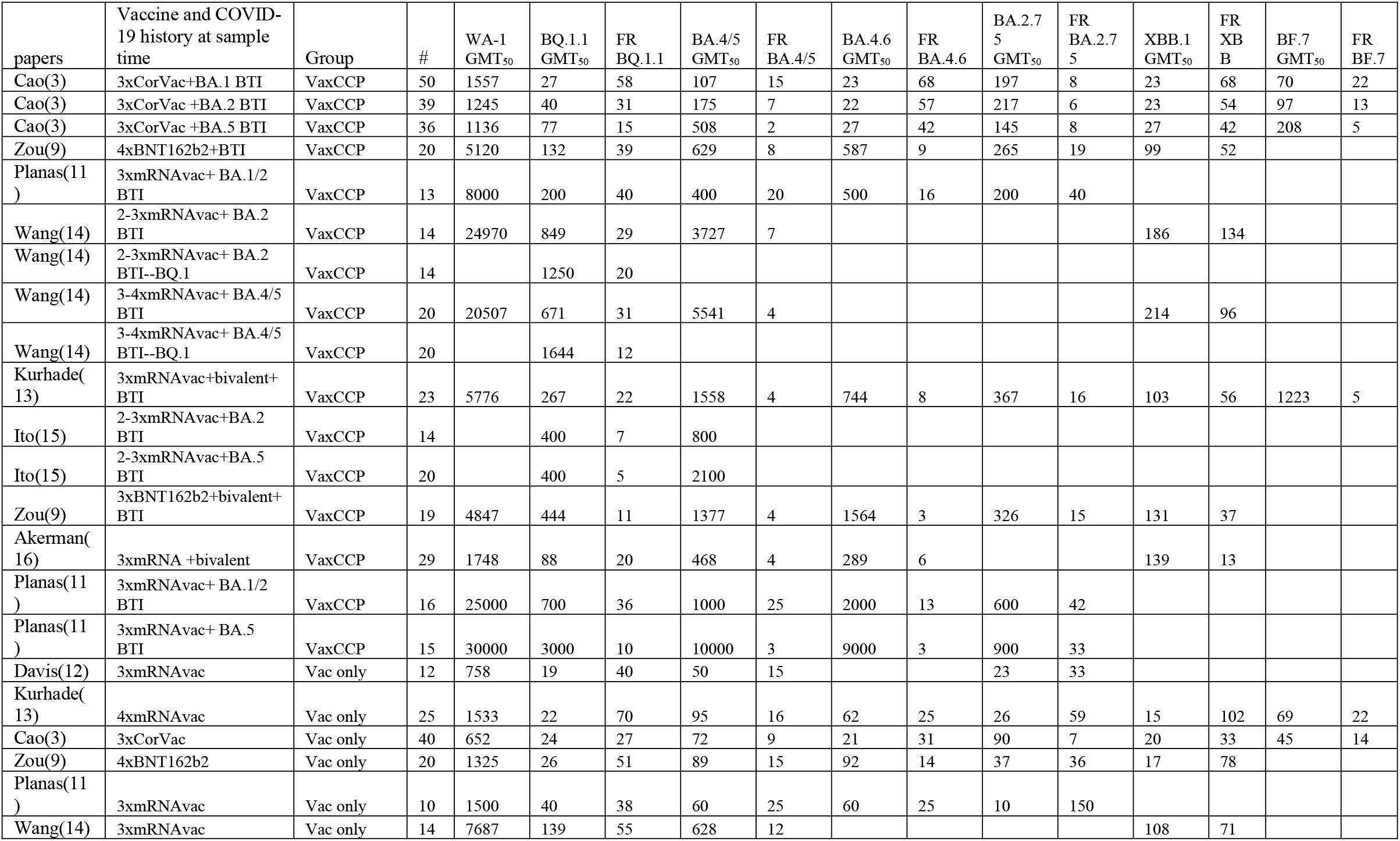
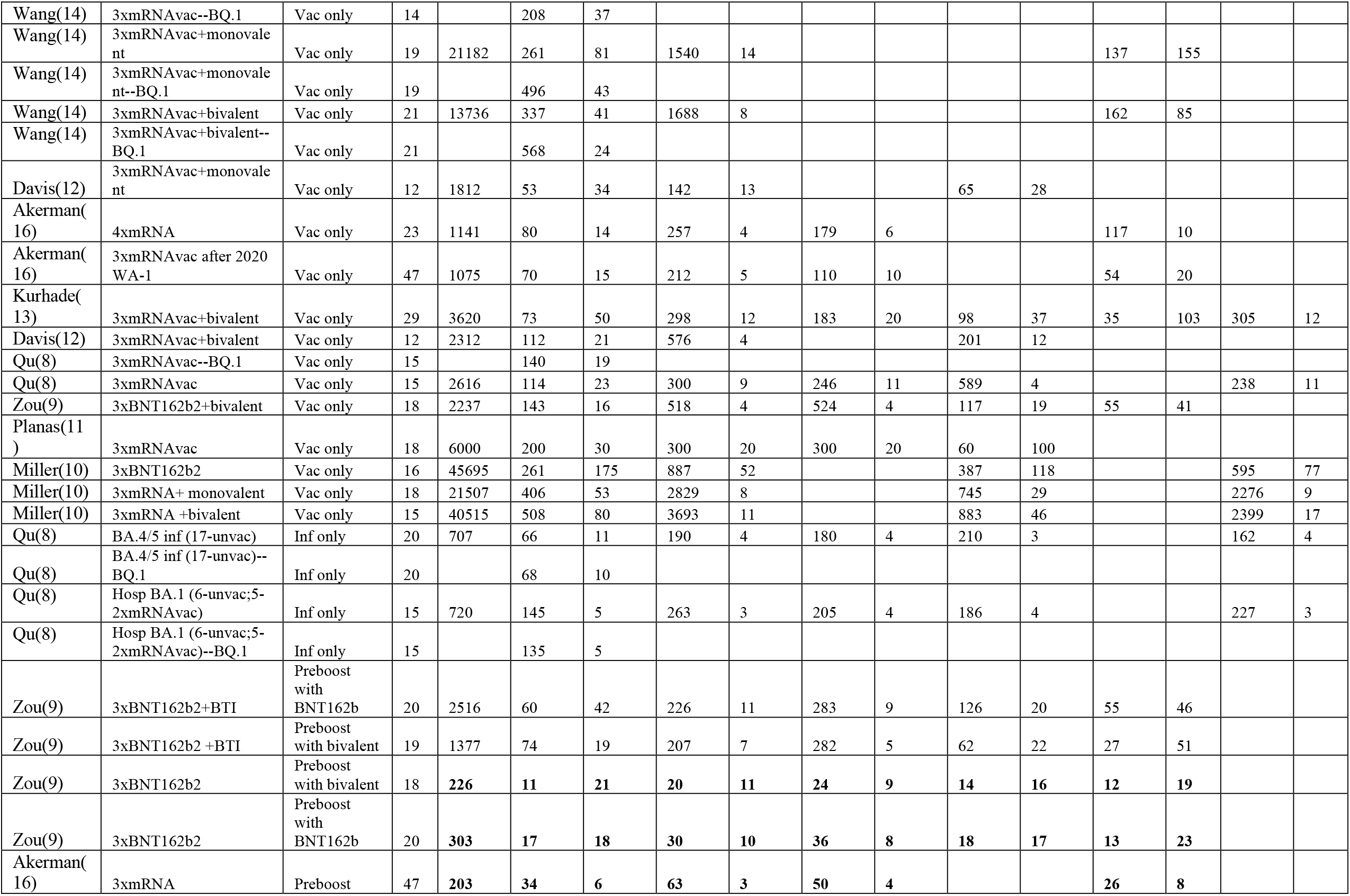
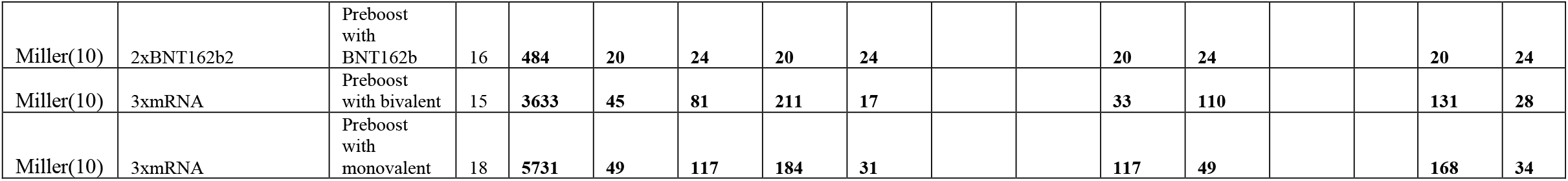
GMT_50_ of different plasma sources against BQ.1.1, BA.4/5, BA.4.6, BA.2.75, XBB.1 and BF.7 and fold-reductions (FR) compared to WA-1.

| papers | Vaccine and COVID-19 history at sample time | Group | # | WA-1 GMT<sub>50</sub> | BQ.1.1 GMT<sub>50</sub> | FR BQ.1.1 | BA.4/5 GMT<sub>50</sub> | FR BA.4/5 | BA.4.6 GMT<sub>50</sub> | FR BA.4.6 | BA.2.7 5 GMT<sub>50</sub> | FR BA.2.7 5 | XBB.1 GMT<sub>50</sub> | FR XB B | BF.7 GMT<sub>50</sub> | FR BF.7 |
| --- | --- | --- | --- | --- | --- | --- | --- | --- | --- | --- | --- | --- | --- | --- | --- | --- |
| Cao(3) | 3xCorVac+BA.1 BTI | VaxCCP | 50 | 1557 | 27 | 58 | 107 | 15 | 23 | 68 | 197 | 8 | 23 | 68 | 70 | 22 |
| Cao(3) | 3xCorVac +BA.2 BTI | VaxCCP | 39 | 1245 | 40 | 31 | 175 | 7 | 22 | 57 | 217 | 6 | 23 | 54 | 97 | 13 |
| Cao(3) | 3xCorVac +BA.5 BTI | VaxCCP | 36 | 1136 | 77 | 15 | 508 | 2 | 27 | 42 | 145 | 8 | 27 | 42 | 208 | 5 |
| Zou(9) | 4xBNT162b2+BTI | VaxCCP | 20 | 5120 | 132 | 39 | 629 | 8 | 587 | 9 | 265 | 19 | 99 | 52 |  |  |
| Planas(11 ) | 3xmRNAvac+ BA.1/2 BTI | VaxCCP | 13 | 8000 | 200 | 40 | 400 | 20 | 500 | 16 | 200 | 40 |  |  |  |  |
| Wang(14) | 2-3xmRNAvac+ BA.2 BTI | VaxCCP | 14 | 24970 | 849 | 29 | 3727 | 7 |  |  |  |  | 186 | 134 |  |  |
| Wang(14) | 2-3xmRNAvac+ BA.2 BTI--BQ.1 | VaxCCP | 14 |  | 1250 | 20 |  |  |  |  |  |  |  |  |  |  |
| Wang(14) | 3-4xmRNAvac+ BA.4/5 BTI | VaxCCP | 20 | 20507 | 671 | 31 | 5541 | 4 |  |  |  |  | 214 | 96 |  |  |
| Wang(14) | 3-4xmRNAvac+ BA.4/5 BTI--BQ.1 | VaxCCP | 20 |  | 1644 | 12 |  |  |  |  |  |  |  |  |  |  |
| Kurhade(13) | 3xmRNAvac+bivalent+ BTI | VaxCCP | 23 | 5776 | 267 | 22 | 1558 | 4 | 744 | 8 | 367 | 16 | 103 | 56 | 1223 | 5 |
| Ito(15) | 2-3xmRNAvac+BA.2 BTI | VaxCCP | 14 |  | 400 | 7 | 800 |  |  |  |  |  |  |  |  |  |
| Ito(15) | 2-3xmRNAvac+BA.5 BTI | VaxCCP | 20 |  | 400 | 5 | 2100 |  |  |  |  |  |  |  |  |  |
| Zou(9) | 3xBNT162b2+bivalent+ BTI | VaxCCP | 19 | 4847 | 444 | 11 | 1377 | 4 | 1564 | 3 | 326 | 15 | 131 | 37 |  |  |
| Akerman(16) | 3xmRNA +bivalent | VaxCCP | 29 | 1748 | 88 | 20 | 468 | 4 | 289 | 6 |  |  | 139 | 13 |  |  |
| Planas(11 ) | 3xmRNAvac+ BA.1/2 BTI | VaxCCP | 16 | 25000 | 700 | 36 | 1000 | 25 | 2000 | 13 | 600 | 42 |  |  |  |  |
| Planas(11 ) | 3xmRNAvac+ BA.5 BTI | VaxCCP | 15 | 30000 | 3000 | 10 | 10000 | 3 | 9000 | 3 | 900 | 33 |  |  |  |  |
| Davis(12) | 3xmRNAvac | Vac only | 12 | 758 | 19 | 40 | 50 | 15 |  |  | 23 | 33 |  |  |  |  |
| Kurhade(13) | 4xmRNAvac | Vac only | 25 | 1533 | 22 | 70 | 95 | 16 | 62 | 25 | 26 | 59 | 15 | 102 | 69 | 22 |
| Cao(3) | 3xCorVac | Vac only | 40 | 652 | 24 | 27 | 72 | 9 | 21 | 31 | 90 | 7 | 20 | 33 | 45 | 14 |
| Zou(9) | 4xBNT162b2 | Vac only | 20 | 1325 | 26 | 51 | 89 | 15 | 92 | 14 | 37 | 36 | 17 | 78 |  |  |
| Planas(11 ) | 3xmRNAvac | Vac only | 10 | 1500 | 40 | 38 | 60 | 25 | 60 | 25 | 10 | 150 |  |  |  |  |
| Wang(14) | 3xmRNAvac | Vac only | 14 | 7687 | 139 | 55 | 628 | 12 |  |  |  |  | 108 | 71 |  |  |
| Wang(14) | 3xmRNAvac--BQ.1 | Vac only | 14 |  | 208 | 37 |  |  |  |  |  |  |  |  |  |  |
| Wang(14) | 3xmRNAvac+monovalent | Vac only | 19 | 21182 | 261 | 81 | 1540 | 14 |  |  |  |  | 137 | 155 |  |  |
| Wang(14) | 3xmRNAvac+monovalent--BQ.1 | Vac only | 19 |  | 496 | 43 |  |  |  |  |  |  |  |  |  |  |
| Wang(14) | 3xmRNAvac+bivalent | Vac only | 21 | 13736 | 337 | 41 | 1688 | 8 |  |  |  |  | 162 | 85 |  |  |
| Wang(14) | 3xmRNAvac+bivalent--BQ.1 | Vac only | 21 |  | 568 | 24 |  |  |  |  |  |  |  |  |  |  |
| Davis(12) | 3xmRNAvac+monovalent | Vac only | 12 | 1812 | 53 | 34 | 142 | 13 |  |  | 65 | 28 |  |  |  |  |
| Akerman(16) | 4xmRNA | Vac only | 23 | 1141 | 80 | 14 | 257 | 4 | 179 | 6 |  |  | 117 | 10 |  |  |
| Akerman(16) | 3xmRNAvac after 2020 WA-1 | Vac only | 47 | 1075 | 70 | 15 | 212 | 5 | 110 | 10 |  |  | 54 | 20 |  |  |
| Kurhade(13) | 3xmRNAvac+bivalent | Vac only | 29 | 3620 | 73 | 50 | 298 | 12 | 183 | 20 | 98 | 37 | 35 | 103 | 305 | 12 |
| Davis(12) | 3xmRNAvac+bivalent | Vac only | 12 | 2312 | 112 | 21 | 576 | 4 |  |  | 201 | 12 |  |  |  |  |
| Qu(8) | 3xmRNAvac--BQ.1 | Vac only | 15 |  | 140 | 19 |  |  |  |  |  |  |  |  |  |  |
| Qu(8) | 3xmRNAvac | Vac only | 15 | 2616 | 114 | 23 | 300 | 9 | 246 | 11 | 589 | 4 |  |  | 238 | 11 |
| Zou(9) | 3xBNT162b2+bivalent | Vac only | 18 | 2237 | 143 | 16 | 518 | 4 | 524 | 4 | 117 | 19 | 55 | 41 |  |  |
| Planas(11) | 3xmRNAvac | Vac only | 18 | 6000 | 200 | 30 | 300 | 20 | 300 | 20 | 60 | 100 |  |  |  |  |
| Miller(10) | 3xBNT162b2 | Vac only | 16 | 45695 | 261 | 175 | 887 | 52 |  |  | 387 | 118 |  |  | 595 | 77 |
| Miller(10) | 3xmRNA+ monovalent | Vac only | 18 | 21507 | 406 | 53 | 2829 | 8 |  |  | 745 | 29 |  |  | 2276 | 9 |
| Miller(10) | 3xmRNA +bivalent | Vac only | 15 | 40515 | 508 | 80 | 3693 | 11 |  |  | 883 | 46 |  |  | 2399 | 17 |
| Qu(8) | BA.4/5 inf (17-unvac) | Inf only | 20 | 707 | 66 | 11 | 190 | 4 | 180 | 4 | 210 | 3 |  |  | 162 | 4 |
| Qu(8) | BA.4/5 inf (17-unvac)--BQ.1 | Inf only | 20 |  | 68 | 10 |  |  |  |  |  |  |  |  |  |  |
| Qu(8) | Hosp BA.1 (6-unvac;5-2xmRNAvac) | Inf only | 15 | 720 | 145 | 5 | 263 | 3 | 205 | 4 | 186 | 4 |  |  | 227 | 3 |
| Qu(8) | Hosp BA.1 (6-unvac;5-2xmRNAvac)--BQ.1 | Inf only | 15 |  | 135 | 5 |  |  |  |  |  |  |  |  |  |  |
| Zou(9) | 3xBNT162b2+BTI | Preboost with BNT162b | 20 | 2516 | 60 | 42 | 226 | 11 | 283 | 9 | 126 | 20 | 55 | 46 |  |  |
| Zou(9) | 3xBNT162b2 +BTI | Preboost with bivalent | 19 | 1377 | 74 | 19 | 207 | 7 | 282 | 5 | 62 | 22 | 27 | 51 |  |  |
| Zou(9) | 3xBNT162b2 | Preboost with bivalent | 18 | **226** | **11** | **21** | **20** | **11** | **24** | **9** | **14** | **16** | **12** | **19** |  |  |
| Zou(9) | 3xBNT162b2 | Preboost with BNT162b | 20 | **303** | **17** | **18** | **30** | **10** | **36** | **8** | **18** | **17** | **13** | **23** |  |  |
| Akerman(16) | 3xmRNA | Preboost | 47 | **203** | **34** | **6** | **63** | **3** | **50** | **4** |  |  | **26** | **8** |  |  |
| Miller(10) | 2xBNT162b2 | Preboost<br>with<br>BNT162b | 16 | **484** | **20** | **24** | **20** | **24** |  |  | **20** | **24** |  |  | **20** | **24** |
| Miller(10) | 3xmRNA | Preboost<br>with bivalent | 15 | **3633** | **45** | **81** | **211** | **17** |  |  | **33** | **110** |  |  | **131** | **28** |
| Miller(10) | 3xmRNA | Preboost<br>with<br>monovalent | 18 | **5731** | **49** | **117** | **184** | **31** |  |  | **117** | **49** |  |  | **168** | **34** |

**Supplementary Table 3.** Neutralizing activity numbers (#) by study cohort.

| papers | Vaccine and COVID-19 history at sample time | groups | WA-1 number | WA-1 neutralizing number | BQ.1.1 number | BQ.1.1 neutralizing number | BA.4/5 number | BA.4/5 neutralizing number | BA.4.6 number | BA.4.6 neutralizing number | BA.2.75 number | BA.2.75 neutralizing number | XBB.1 number | XBB.1 neutralizing number | BF.7 number | BF.7 neutralizing number |
| --- | --- | --- | --- | --- | --- | --- | --- | --- | --- | --- | --- | --- | --- | --- | --- | --- |
| Cao(3) | 3xCorVac+BA.1 BTI | VaxCCP | 50 | 50 |  |  | 50 | 50 |  |  | 50 | 50 |  |  | 50 | 49 |
| Cao(3) | 3xCorVac +BA.2 BTI | VaxCCP | 39 | 39 |  |  | 39 | 39 |  |  | 39 | 39 |  |  | 39 | 38 |
| Cao(3) | 3xCorVac +BA.5 BTI | VaxCCP | 36 | 36 |  |  | 36 | 36 |  |  | 36 | 36 |  |  | 36 | 36 |
| Zou(9) | 4xBNT162b2+BTI | VaxCCP | 20 | 20 | 20 | 20 | 20 | 20 | 20 | 20 | 20 | 20 | 20 | 19 |  |  |
| Planas(11) | 3xmRNAvac+ BA.1/2 BTI | VaxCCP | 13 | 13 | 13 | 11 | 13 | 12 | 13 | 13 | 13 | 11 |  |  |  |  |
| Wang(14) | 2-3xmRNAvac+ BA.2 BTI | VaxCCP | 14 | 14 | 14 | 13 | 14 | 14 |  |  |  |  | 14 | 8 |  |  |
| Wang(14) | 2-3xmRNAvac+ BA.2 BTI--BQ.1 | VaxCCP |  |  | 14 | 13 |  |  |  |  |  |  |  |  |  |  |
| Wang(14) | 3-4xmRNAvac+ BA.4/5 BTI | VaxCCP | 20 | 20 | 20 | 18 | 20 | 20 |  |  |  |  | 20 | 14 |  |  |
| Wang(14) | 3-4xmRNAvac+ BA.4/5 BTI--BQ.1 | VaxCCP |  |  | 20 | 20 |  |  |  |  |  |  |  |  |  |  |
| Kurhade(13) | 3xmRNAvac+bivalent+BTI | VaxCCP | 23 | 23 | 23 | 22 | 23 | 23 | 23 | 23 | 23 | 23 | 23 | 22 | 23 | 23 |
| Ito(15) | 2-3xmRNAvac+BA.2 BTI | VaxCCP |  |  | 14 | 11 | 14 | 13 |  |  |  |  |  |  |  |  |
| Ito(15) | 2-3xmRNAvac+BA.5 BTI | VaxCCP |  |  | 20 | 14 | 20 | 20 |  |  |  |  |  |  |  |  |
| Zou(9) | 3xBNT162b2+bivalent+BTI | VaxCCP | 19 | 19 | 19 | 19 | 19 | 19 | 19 | 19 | 19 | 20 | 19 | 19 |  |  |
| Akerman(16) | 3xmRNA +bivalent | VaxCCP | 29 | 20 | 29 | 29 | 29 | 29 | 29 | 29 |  |  | 29 | 29 |  |  |
| Planas(11) | 3xmRNAvac+ BA.1/2 BTI | VaxCCP | 16 | 16 | 16 | 16 | 16 | 16 | 16 | 16 | 16 | 16 |  |  |  |  |
| Planas(11) | 3xmRNAvac+ BA.5 BTI | VaxCCP | 15 | 15 | 15 | 15 | 15 | 15 | 15 | 15 | 15 | 15 |  |  |  |  |
| Davis(12) | 3xmRNAvac | Vac only | 12 | 12 | 12 | 6 | 12 | 11 |  |  | 12 | 9 |  |  |  |  |
| Kurhade(13) | 4xmRNAvac | Vac only | 25 | 25 | 25 | 15 | 25 | 23 | 25 | 22 | 25 | 17 | 25 | 8 | 25 | 21 |
| Cao(3) | 3xCorVac | Vac only | 40 | 40 |  |  | 40 | 39 |  |  | 40 | 40 |  |  | 40 | 37 |
| Zou(9) | 4xBNT162b2 | Vac only | 20 | 20 | 20 | 15 | 20 | 8 | 20 | 19 | 20 | 19 | 20 | 10 |  |  |
| Planas(11) | 3xmRNAvac | Vac only | 10 | 10 | 10 | 6 | 10 | 7 | 10 | 5 | 10 | 3 |  |  |  |  |
| Wang(14) | 3xmRNAvac | Vac only | 14 | 14 | 14 | 5 | 14 | 14 |  |  |  |  | 14 | 3 |  |  |
| Wang(14) | 3xmRNAvac--BQ.1 | Vac only |  |  | 14 | 6 |  |  |  |  |  |  |  |  |  |  |
| Wang(14) | 3xmRNAvac+monovalent | Vac only | 19 | 19 | 19 | 15 | 19 | 19 |  |  |  |  | 19 | 8 |  |  |
| Wang(14) | 3xmRNAvac+monovalent--BQ.1 | Vac only |  |  | 19 | 16 |  |  |  |  |  |  |  |  |  |  |
| Wang(14) | 3xmRNAvac+bivalent | Vac only | 21 | 21 | 21 | 16 | 21 | 21 |  |  |  |  | 21 | 9 |  |  |
| Wang(14) | 3xmRNAvac+bivalent--BQ.1 | Vac only |  |  | 21 | 18 |  |  |  |  |  |  |  |  |  |  |
| Davis(12) | 3xmRNAvac+monovalent | Vac only | 12 | 12 | 12 | 9 | 12 | 12 |  |  | 12 | 10 |  |  |  |  |
| Akerman(16) | 4xmRNA | Vac only | 23 | 23 | 23 | 20 | 23 | 23 | 23 | 22 |  |  | 23 | 17 |  |  |
| Akerman(16) | 3xmRNAvac after 2020 WA-1 | Vac only | 47 | 46 | 47 | 34 | 47 | 45 | 47 | 43 |  |  | 47 | 32 |  |  |
| Kurhade(13) | 3xmRNAvac+bivalent | Vac only | 29 | 29 | 29 | 26 | 29 | 29 | 29 | 28 | 29 | 28 | 29 | 20 | 29 | 28 |
| Davis(12) | 3xmRNAvac+bivalent | Vac only | 12 | 12 | 12 | 10 | 12 | 12 |  |  | 12 | 11 |  |  |  |  |
| Qu(8) | 3xmRNAvac--BQ.1 | Vac only |  |  | 15 | 14 |  |  |  |  |  |  |  |  |  |  |
| Qu(8) | 3xmRNAvac | Vac only | 15 | 15 | 15 | 12 | 15 | 15 | 15 | 15 | 15 | 14 |  |  | 15 | 14 |
| Zou(9) | 3xBNT162b2+bivalent | Vac only | 18 | 18 | 19 | 18 | 18 | 18 | 19 | 19 | 19 | 18 | 19 | 18 |  |  |
| Planas(11) | 3xmRNAvac | Vac only | 18 | 18 | 18 | 16 | 18 | 18 | 18 | 18 | 18 | 13 |  |  |  |  |
| Miller(10) | 3xBNT162b2 | Vac only | 16 | 16 | 16 | 16 | 16 | 16 |  |  | 16 | 16 |  |  | 16 | 16 |
| Miller(10) | 3xmRNA+monovalent | Vac only | 18 | 18 | 18 | 18 | 18 | 18 |  |  | 18 | 18 |  |  | 18 | 18 |
| Miller(10) | 3xmRNA +bivalent | Vac only | 15 | 15 | 15 | 15 | 15 | 15 |  |  | 15 | 15 |  |  | 15 | 15 |
| Qu(8) | BA.4/5 inf (17-unvac) | Inf only | 20 | 20 | 20 | 12 | 20 | 15 | 20 | 14 | 20 | 15 |  |  | 20 | 14 |
| Qu(8) | BA.4/5 inf (17-unvac)--BQ.1 | Inf only |  |  | 20 | 11 |  |  |  |  |  |  |  |  |  |  |
| Qu(8) | Hosp BA.1 (6-unvac;5-2xmRNAvac) | Inf only | 15 | 14 | 15 | 13 | 15 | 11 | 15 | 12 | 15 | 8 |  |  | 15 | 14 |
| Qu(8) | Hosp BA.1 (6-unvac;5-2xmRNAvac)--BQ.1 | Inf only |  |  | 15 | 14 |  |  |  |  |  |  |  |  |  |  |
| Zou(9) | 3xBNT162b2+BTI | Preboost with BNT162b | 20 | 20 | 20 | 16 | 20 | 19 | 20 | 19 | 20 | 17 | 20 | 17 |  |  |
| Zou(9) | 3xBNT162b2 +BTI | Preboost with bivalent | 19 | 19 | 19 | 16 | 19 | 19 | 19 | 19 | 19 | 17 | 19 | 12 |  |  |
| Zou(9) | **3xBNT162b2** | **Preboost with bivalent** | **18** | **18** | **19** | **1** | **18** | **9** | **19** | **9** | **19** | **5** | **19** | **2** |  |  |
| Zou(9) | **3xBNT162b2** | **Preboost with BNT162b** | **20** | **19** | **20** | **6** | **20** | **12** | **20** | **12** | **20** | **8** | **20** | **5** |  |  |
| Akerman (16) | **3xmRNA** | **Preboost** | **23** | **23** | **23** | **16** | **23** | **21** | **23** | **19** |  |  | **23** | **14** |  |  |
| Miller(10) | **2xBNT162b2** | **Preboost with BNT162b** | **16** | **16** | **16** | **6** | **16** | **5** |  |  | **16** | **7** |  |  | **16** | **4** |
| Miller(10) | **3xmRNA** | **Preboost with bivalent** | **15** | **15** | **15** | **13** | **15** | **15** |  |  | **15** | **9** |  |  | **15** | **14** |
| Miller(10) | **3xmRNA** | **Preboost with monovalent** | **18** | **18** | **18** | **15** | **18** | **18** |  |  | **18** | **18** |  |  | **18** | **17** |

**Supplementary Figure 1.**
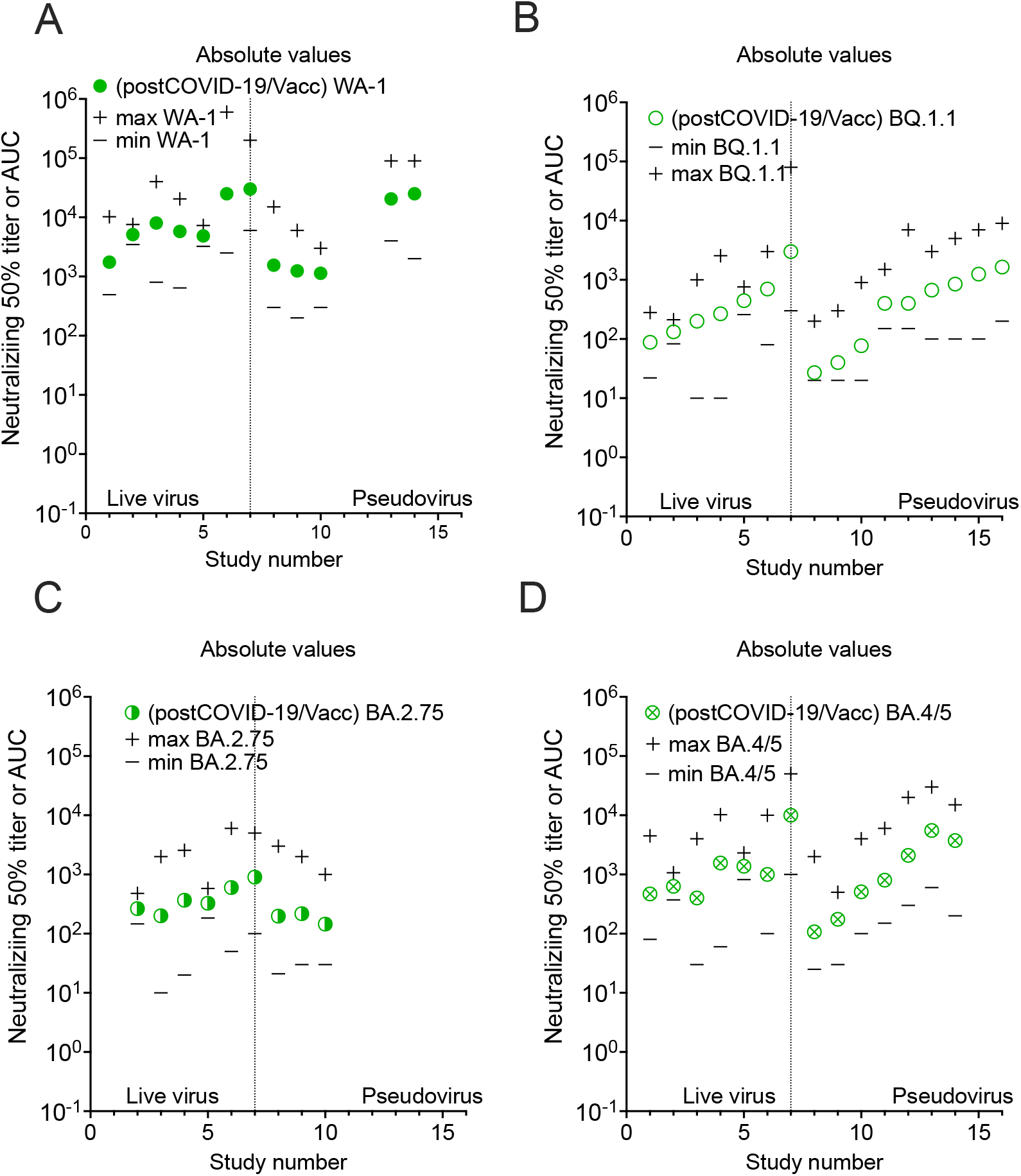
Plasma GMT_50_ from post boosted vaccinations and COVID-19 sorted by study cohort with live virus assays on the left and pseudovirus on right, with individual sample minimum and maximum dilution titer also shown. A) WA-1; B) BQ.1.1; C) BA.2.75; D) BA.4/5.

**Supplementary Figure 2.**
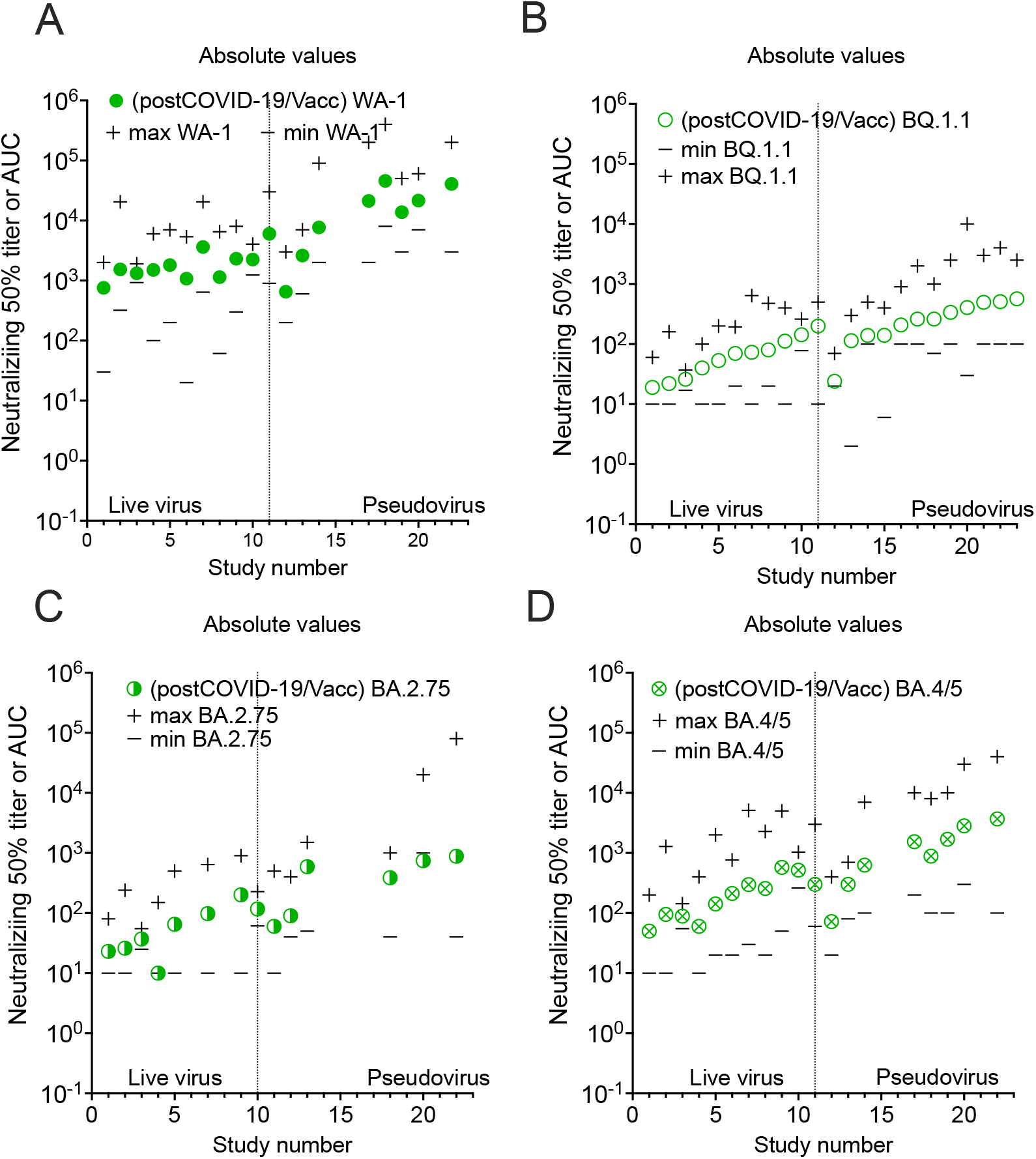
Plasma GMT_50_ from boosted vaccinations only without COVID-19 sorted by study cohort with live virus assays on the left and pseudovirus on right with with individual sample minimum and maximum dilution titer also shown. A) WA-1; B) BQ.1.1; C) BA.2.75; and D) BA.4/5.

**Supplementary Figure 3.**
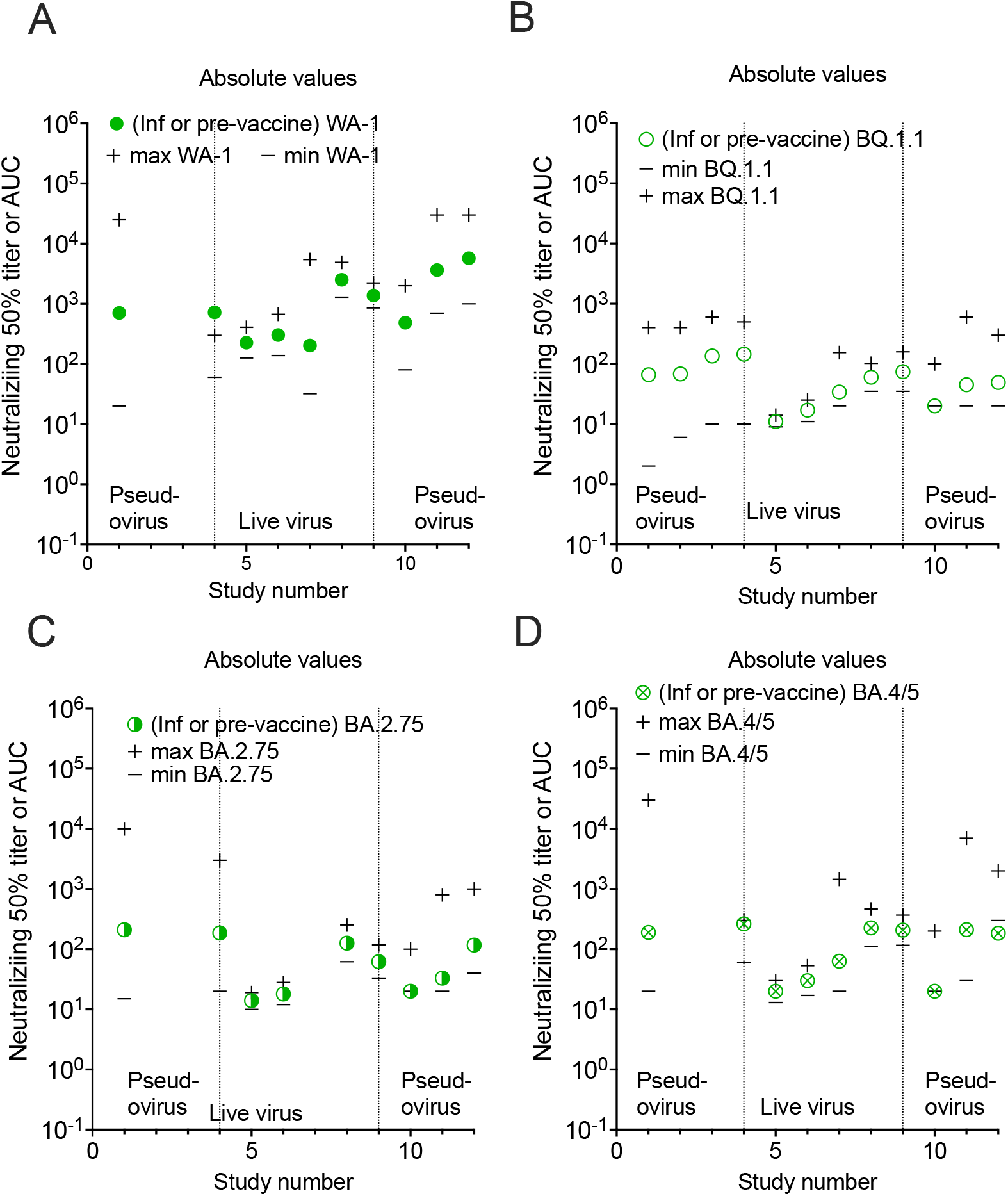
Plasma GMT_50_ from Omicron infection alone and also pre-boosted in 2021 or 2022 6 to 11 months after last vaccine dose sampled sorted by study cohort with live virus assays on the left and pseudovirus on right with minimum and maximum dilution titer also shown. A) WA-1; B) BQ.1.1; C) BA.2.75; and D) BA.4/5.

